# Causal variant underestimation is a major overlooked driver of sequence-to-function model underperformance

**DOI:** 10.64898/2026.09.08.750172

**Authors:** Shiron Drusinsky, Katherine S. Pollard

## Abstract

Deep learning sequence-to-function (S2F) models represent tremendous promise for functionally fine-mapping causal variants associated with traits and disease. Yet it remains unclear precisely how effective they are at this task. Generally, S2F models perform well at classifying putatively causal expression quantitative trait loci (eQTL) SNVs, yet dramatically underperform linear baselines at ranking different individuals’ gene expression values from their whole genome sequence, which directly calls into question their ability to fine-map a locus by properly weighting the effects of variants onto gene expression. Here, using the state-of-the-art S2F model AlphaGenome, we systematically compared predicted effect sizes for fine-mapped eQTLs to nearby putatively non-causal SNVs, finding that AlphaGenome pervasively underestimates the effects of most causal variants and fails to successfully fine-map eQTLs in most loci for this reason. We show that misdirected variant effect predictions and negative cross-individual correlations, widely cited as major challenges facing S2F expression modeling, are not egregious errors but a chance consequence of weak, noisy attributions when causal variants are underestimated. Our results suggest that a failure to detect local variant effects onto enhancer activity is a cause of underestimation. Additionally, we find that underestimated variants are enriched far from their target gene, suggesting inadequate enhancer-gene linking causes underestimation even when local effects are well-detected. Finally, our results clarify when S2F models are reliable for fine-mapping objectives: while they pervasively underestimate most causal variants, demonstrating limited fine-mapping utility for most loci, the top *∼*0.1% most prominent variant effect predictions are strongly enriched for causal variants with directionally correct predictions that are consistent across model replicates, suggesting extremely strong predictions are broadly trustworthy. Our results demonstrate that causal variant underestimation is a core issue facing S2F expression predictors, with future improvements dependent on better local activity detection and enhancer-gene linking.

## 1 Introduction

Sequence-to-function (S2F) models take DNA sequence as input and predict outcomes of thousands of context specific functional genomics experiments – including DNA accessibility, transcription factor binding, and gene expression assays – in unseen loci with high accuracy, suggesting tremendous promise for decoding the “grammar” underlying the noncoding genome [1–7]. There is therefore great interest in applying these models to functionally fine-map causal variants in trait- and disease-associated loci based on their predicted perturbations to gene regulation. Indeed, S2F model predictions have been shown to improve causal variant prioritization when combined with statistical fine-mapping approaches [8, 9].

Yet it remains unclear precisely how adept S2F models are at fine-mapping loci on their own, because standard evaluations are poorly suited to evaluate this task. Typical benchmarks compare predicted versus observed effects of fine-mapped expression quantitative trait loci (fm-eQTLs) and evaluate the ability of S2F models to correctly rank and predict the direction of effect of these variants. Other benchmarks ask models to classify fm-eQTLs relative to matched, putatively non-causal variants based on the predicted effect sizes of each, but these matched controls typically come from different loci [1–3, 6, 10]. Though S2F models – including AlphaGenome[1], the current state-of-the-art – are highly performant on these tasks, these evaluation strategies ultimately fail to establish the extent to which S2F models prominently upweight causal variants above nearby non-causal ones, as is needed to evaluate their fine-mapping capabilities.

More recently, various groups have demonstrated that S2F models dramatically underperform linear baselines at predicting gene expression differences between individuals given their personal whole genome sequences [11–15]. This implies improper variant effect predictions, and seemingly contradicts promising results from standard fm-eQTL benchmarks. Independently, we noticed S2F models often predict surprisingly weak effect sizes for most fm-eQTLs (see **Figs. 1A, S1A** for example), despite strong benchmark results, an observation that is also prevalent in the literature [1, 3] – including among higher-confidence fm-eQTLs that replicate in separate cohorts [9] – but has sparked little discussion. Because personal genome predictions aggregate all variants in a window, accuracy requires that models upweight causal variants while downweighting the many surrounding non-causal ones. We therefore hypothesize these seemingly disparate observations are closely related, and reflect poor S2F model fine-mapping capabilities at most loci, with causal variants often receiving weaker than expected attributions relative to surrounding background variants.

**Figure 1:**
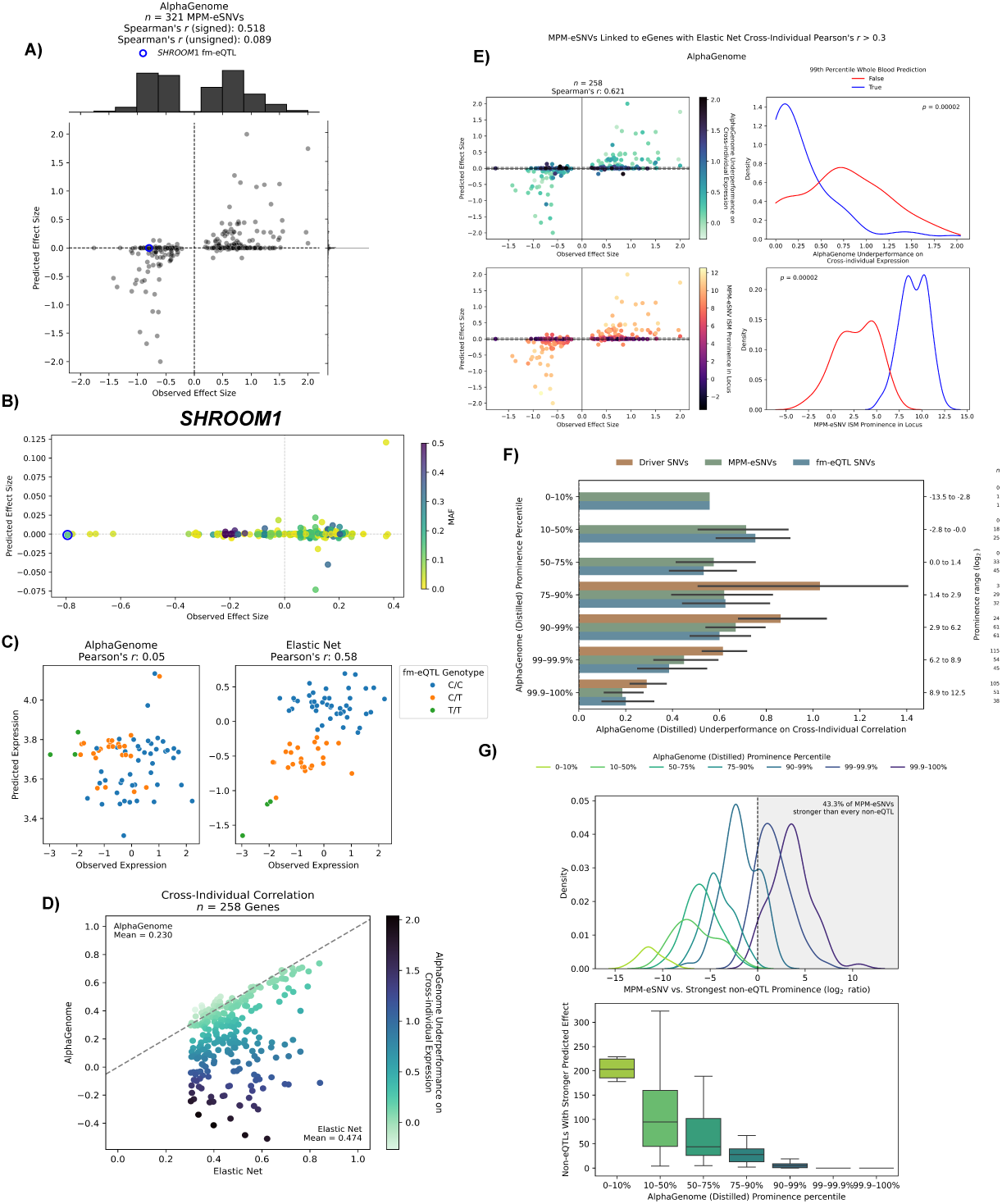
AlphaGenome pervasively underestimates expression effects of causal variants, leading to fine-mapping issues. **A)** Predicted versus observed effect size (eQTL betas) for maximum predicted magnitude eSNVs (MPM-eSNVs). Predicted effect sizes are concentrated around 0 despite strong signed correlation across SNVs. Strong wrong directional predictions are rare. **B)** Predicted versus observed effect size for all observed SNVs within 131kb of *SHROOM1*. A fm-eQTL is annotated in a blue circle, and its predicted effect size is indistinguishable from background SNVs, a fine-mapping failure. **C)** The fine-mapping failure from (B) causes AlphaGenome to underperform a linear baseline on predicting gene expression differences across individuals using their personal whole genome sequences. The elastic net properly differentiates individuals based on their genotype of the fm-eQTL from (B); AlphaGenome does not. Each point is a different individual and their observed and predicted expression of *SHROOM1*. **D)** Cross-individual correlations for many genes (as in panel C) from AlphaGenome (Y-axis) and the elastic net baseline (X axis). Points are colored by a scaled underperformance metric (**Methods**). AlphaGenome underperforms for most genes, suggesting widespread issues with S2F fine-mapping capabilities. **E)** Predicted versus observed effect sizes for MPM-eSNVs colored by AlphaGenome’s personal genome underperformance on these SNVs’ eGenes (top) and their prominence relative to nearby SNVs (bottom). Dashed lines represent the global 99th percentile prediction threshold, used to coarsely stratify SNVs with a predicted effect size near zero from the rest, and to form the two groups on the right. P-values are from a permutation test. Personal genome prediction accuracy is associated with MPM-eSNV prominence. **F)** AlphaGenome underperformance on the personal genome task for eGenes linked to variants with different prominence levels. Binned prominence rankings appear on the left; underlying prominence ranges appear on the right, along with the number of genes in each group. When multiple variants from a class are linked to the same eGene, their prominence is averaged. **G)** MPM-eSNVs are binned into different prominence quantiles (color) and their prominence is compared to that of the strongest non-eQTL within the same window (top). 43.3% are stronger than nearby non-eQTLs, but many have comparable effects; only the 99.9th percentile prominence predictions stand out from non-eQTLs. The number of non-eQTLs with greater prominence than MPM-eSNVs (bottom). The remaining 56.7% of MPM-eSNVs are weaker than many nearby non-eQTLs.

To investigate this hypothesis, we systematically compared predicted effects of fm-eQTLs to those of neighboring SNVs using data from the Genotype-Tissue Expression (GTEx) consortium [16]. Our study clarifies that S2F models poorly fine-map most loci due to pervasive causal variant underestimation. We identify likely causes of underestimation, delineate when variant effect predictions can and cannot be trusted, and unify several challenges facing S2F expression models under this single overlooked issue.

## 2 Results

### 2.1 Pervasive causal variant underestimation leads to S2F fine-mapping challenges

To analyze the accuracy of current S2F models on regulatory variant effect predictions, we focus on AlphaGenome [1]. Using 131kb transcription-start site (TSS)-centered sequences containing fine-mapped (fm; PIP *>* 0.9) ‘Whole Blood’ eQTLs from GTEx [16], we found that most fm-eQTL SNVs were concentrated near a predicted effect size of 0. The aggregate performance metric was nonetheless strong (signed Spearman’s *r* = 0.45), but it was predominantly driven by a subset of points with high leverage (**Fig. S1A**), rather than a consistent positive relationship between predicted and observed effects. Similar patterns of reasonable across-fm-eQTL correlations, despite widespread underestimation, can be found in benchmarking figures from S2F model publications (see Figs. 4D and Extended Data Fig. 5H in [1], Fig. S9B in [3], and Fig. 1D and Extended Data Figs. 1B-D in [9]). This suggests that underestimation is common over an entire class of S2F models, and is not a technical artifact from our methodology, but is a finding that has – to our knowledge – gone mostly overlooked in discussion within the literature thus far. Pervasive underestimation of causal SNVs is important, because it could imply issues with fine-mapping, variant interpretation, and enhancer-gene linking for many causal regulatory loci. This motivated us to further investigate this observation.

To verify that AlphaGenome was not assigning the causal source of the eQTL association to other plausible SNV candidates instead, we replaced fm-eQTL SNVs with other SNVs in tight Linkage Disequilibrium (LD; *R*^2^ *>* 0.9) if those SNVs were predicted by AlphaGenome to cause a larger magnitude change in expression of the same gene than the fm-eQTL SNV. We refer to these as maximum predicted magnitude eSNVs (MPM-eSNVs). Few fm-eQTL SNVs had variants in tight LD, so the MPM-eSNV set differs by only 36/321 SNV-eGene pairs. Accordingly, AlphaGenome continued to severely underestimate MPM-eSNVs (**Fig. 1A**). We repeated this experiment using MPM-eSNVs in an expanded 1Mb input sequence window, again obtaining nearly identical results (**Fig. S1B**), confirming AlphaGenome is also not upweighting other candidate linked SNVs outside the original 131kb window instead of fm-eQTL SNVs. Throughout the rest of this manuscript, we therefore use 131kb sequences for computational feasibility, and we use MPM-eSNVs to rule out the possibility of disagreement between AlphaGenome and statistical fine-mapping as an explanation for our findings.

Small predicted effect sizes are not, on their own, problematic because S2F models are not necessarily expected to predict effect sizes on the same scale as eQTL tests. However, tasks such as fine-mapping require that models at least correctly rank the effects of different variants within a locus onto expression of a given gene, and weight them appropriately relative to one another. To understand whether causal SNVs with weak predicted effects ranked higher than background SNVs, we compared the predicted and observed effect sizes for all SNVs with a reported effect size estimate (eQTL beta) by GTEx – i.e., minor allele frequency (MAF) *≥* 1% – and within 131kb of the *SHROOM1* transcription start site (TSS), a gene with a fm-eQTL SNV (**Fig. 1A**). We found that this fm-eQTL SNV not only had a weak AlphaGenome predicted effect, but its effect was indistinguishable from those of most other SNVs in the locus (**Fig. 1B**). When searching for a different MPM-eSNV, we found AlphaGenome did not upweight another candidate SNV in LD instead (**Fig. S1C**), confirming that AlphaGenome definitively underestimates the *SHROOM1* eQTL and would not prioritize any other SNV that could underlie the expression association. This is consequential; due to its inability to correctly upweight the *SHROOM1* fm-eQTL, a fine-mapping failure, AlphaGenome also drastically underperformed a linear baseline on the task of ranking gene expression across individuals using their personal genome sequence (**Fig. 1C**). This is a well-documented issue facing S2F modeling efforts [11–15], affecting most genes to varying extents (**Fig. 1D**).

To investigate whether AlphaGenome underestimates causal SNVs relative to background SNVs from a broader set of genes, we analyzed its ability to predict Whole Blood gene expression differences across individuals, per gene, given their personal whole genome sequences. This is effectively a fine-mapping task that is well-suited to reflect underestimation, as accurate ranking of gene expression across people requires the model to properly upweight important SNVs and downweight unimportant ones, so accuracy improves when causal SNVs stand out prominently against the weaker and inert SNVs around them, and suffers when they do not – as in the *SHROOM1* case (**Fig. 1B**). We therefore compared AlphaGenome’s personal genome predictions against those of elastic net models, a linear approach commonly used to establish the cross-individual accuracy attainable from common, local genetic variation [11–15, 17]. Matching or exceeding this baseline suggests important SNVs were prominent enough to drive the prediction, while falling short indicates their underestimation relative to nearby background SNVs. To focus on genes where genetic variants are expected to contribute highly to gene expression differences across individuals, and accurate personal genome predictions are therefore possible, we focused our analyses on genes with at least one Whole Blood fm-eQTL and where an elastic net achieves a cross-individual correlation of at least 0.3. This left us with 366 Whole Blood fm-eQTL SNVs and 321 MPM-eSNVs linked to 258 eGenes (**Methods**).

We found that AlphaGenome personal genome predictions were significantly more accurate, and more closely matched elastic net accuracy, when MPM-eSNVs had larger predicted effect magnitudes (**Fig. 1E**, see **Fig. S5** for examples). Conversely, genes with weakly predicted MPM-eSNVs had less accurate personal genome predictions, suggesting these MPM-eSNVs had not just weak absolute predicted effects, but failed to stand out against other background SNVs in the same locus. To confirm, we assessed the prominence of SNV predictions, defined as the log_2_ ratio of the magnitude of a focal SNV’s predicted effect relative to that of the median SNV around the same gene (**Methods**). Indeed, we found that the same MPM-eSNVs with weak predicted effects, and whose eGenes had inaccurate personal genome predictions, also suffered from weak prominence. Conversely, the most prominent SNVs were linked to eGenes with highly accurate personal genome predictions (**Figs. 1E-F**). Further, while 43% of MPM-eSNVs had stronger predicted effects than all nearby common SNVs that lack an association with expression (non-eQTL SNVs; **Methods**), the remaining were typically outranked by many, and often by substantial amounts (**Fig. 1G**). Even among the 43% that did lead, many exceeded the closest non-eQTL SNV by only a narrow margin. Most loci therefore contain other putatively non-causal SNVs with comparable or even stronger cumulative predicted effects, against which causal SNVs fail to prominently stand out (i.e., poor fine-mapping), explaining the inaccurate personal genome predictions (see **Figs. S3-S4** for examples). The root issue is an insufficient gap in attributions between causal and surrounding background SNVs that are expected to be weaker or non-causal; we refer to this as causal SNV underestimation relative to these background SNVs. As AlphaGenome’s personal genome predictions underperformed linear baselines for most genes (**Fig. 1D**), our results suggest that AlphaGenome pervasively underestimates expression effects of most candidate causal SNVs, not just in absolute magnitude but also relative to background SNVs from the same locus, and in many cases severely. This is akin to a high false negative rate in classification settings, and causes fine-mapping failures that result in inaccurate personal genome predictions, a well-documented S2F modeling challenge.

### 2.2 Causal variant underestimation explains other major S2F challenges

We next hypothesized that SNV underestimation may also explain negatively correlated personal genome predictions, where S2F models predict individuals with truly higher expression as having lower expression and vice versa (see **Fig. S3** for a few examples). Wrong directional variant effect predictions have been widely cited as the cause of this concerning phenomenon [11, 13, 14], and therefore are considered a major challenge for S2F models. Yet, among causal SNVs, we observed few instances of strong, wrong directional predictions (**Figs. 1A, S1A**), consistent with benchmarking results from the AlphaGenome and Borzoi manuscripts [1, 3]. Instead, wrong directional causal variant effect predictions were most common among underestimated causal SNVs (**Fig. 2A**), where we found many ordinary non-eQTLs from the same loci carry comparable or stronger predicted effects (**Fig. 1G**), but were extremely rare for the most prominent SNVs.

**Figure 2:**
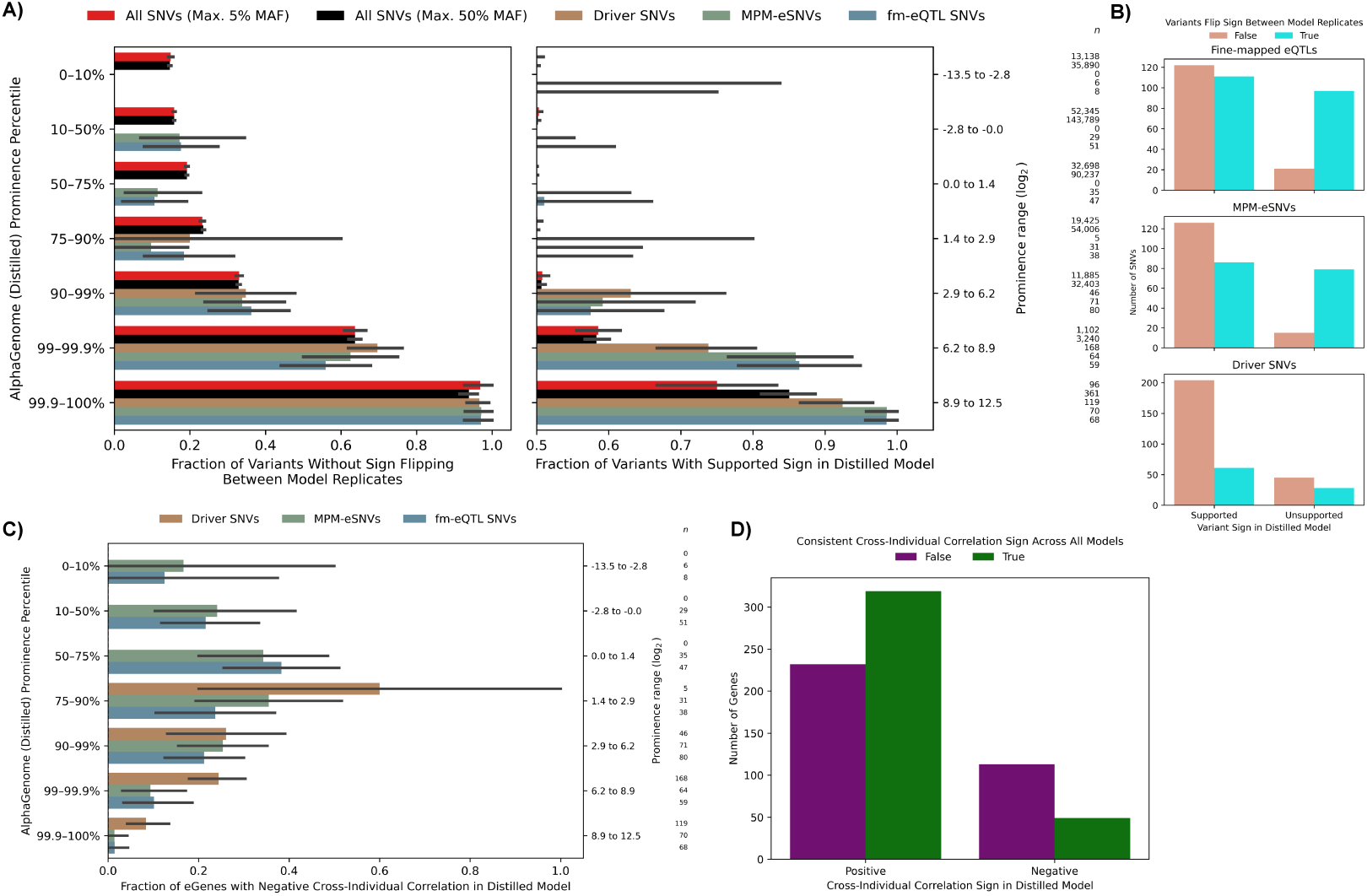
Directional mispredictions and negative cross-individual correlations are byproducts of causal variant underestimation and occur by chance rather than confident model errors. **A)** Prominence of variants from different classes, consistent with Fig. 1F. “All SNVs (Max. 50% MAF)” includes all SNVs our dataset (**Methods**); “All SNVs (Max. 5% MAF)” includes all SNVs under 5% MAF. X-axis shows fraction of variants, stratified by prominence range and variant class, that do not have a predicted direction of effect that flips to the opposite sign in a different cross-validation fold (left), or whose direction of effect by the distilled model is consistent with their eQTL beta sign (right). Sample sizes on the right represent the number of variants from each class in each prominence bin. Wrong directional variant effect predictions are most common for underestimated causal SNVs but rare otherwise. Wrong directional predictions coincide with high model uncertainty. **B)** Variants from different classes (rows) stratified by whether their predicted direction of effect, by the distilled model, is consistent with their eQTL beta (X-axis) and whether that predicted direction flips to the opposite sign when predictions are drawn from a different cross-validation fold (color). **C)** Prominence of variants from different classes, consistent with Fig. 1F. X-axis represents fraction of genes whose cross-individual correlation is negative by the distilled AlphaGenome model. When multiple variants from a class are linked to the same eGene, their prominence is averaged. **D)** Among 162 genes whose cross-individual correlations are negative in the distilled AlphaGenome model (right), 113 revert to positive when predictions are drawn from one of the cross-validation folds (purple), while only 49 do not (green). Among those that were positive in the distilled model (left), 232 revert to negative while 319 stay positive. Cross-individual correlation signs are generally unstable due to lack of a sufficiently prominent variant to anchor the predictions. See Fig. S5 for examples of such anchoring.

To test if these effects were confidently flipped, we examined predictions across four AlphaGenome models trained on independent subsets of training data, and compared them with predictions from the distilled model (used elsewhere throughout this manuscript unless otherwise stated), which was trained on all subsets of training data. Variant effect predictions were highly correlated between the AlphaGenome distilled model and cross-validation replicates, supporting our usage of these replicates to estimate uncertainty in the distilled model (**Fig. S2A**). For most causal SNVs with mispredicted directional effects in the distilled model, the sign flipped back in at least one of these cross-validation fold replicates (**Fig. 2B**), suggesting these are chance sign-flips rather than confident mispredictions. More broadly, variant-effect predictions routinely flipped direction between model folds for all but the most prominent SNVs, suggesting high model uncertainty everywhere but the very top of the dynamic range (99.9th percentile prominence and above), where predicted effects are mostly supported by eQTL betas (**Fig. 2A**). Thus, even supported variant directional effects often flipped to unsupported ones in one of the cross-validation folds (**Fig. 2B**), due to general uncertainty that coincides with underestimation.

Negatively correlated personal genome predictions behaved similarly, as they were more common when causal SNVs were underestimated (**Fig. 2C**) and typically reverting to positive when predictions were drawn from a different model fold (**Fig. 2D**), as expected if they arise from chance sign-flips rather than confident mispredictions that can reliably hold a gene’s ranking of individuals in the wrong direction. For the same reason, even strong, positive cross-individual correlations can be driven by weakly prominent SNVs and occur by chance due to a fortunate combination of variant effect predictions in one model fold that does not hold in others, motivating the use of many model replicates when analyzing personal genome predictions (**Figs. 2D, S4**). Indeed, we observed wide variation in cross-individual correlation across model folds for most genes – including genes that approached but never crossed zero, and genes that matched the elastic net baseline in one fold yet fell well short in another (**Fig. S6**). With only four model folds, the number of genes that flip sign between them is likely undercounted. We verified this wide variation could not be explained by large differences in attributions between the three cross-validation folds that included each gene in its train set, versus the single remaining fold that did not (**Fig. S7**).

We repeated our analyses using driver SNVs, the minimal set of SNVs that best explain AlphaGenome’s personal genome predictions for a given gene due to their large predicted effect sizes and typically common MAFs (**Methods**). We investigated drivers because their directional mispredictions were originally used to explain negatively correlated personal genome predictions, implicating variant directional mispredictions as a major challenge for S2F models. By construction, drivers are among the most prominent SNVs in the locus (**Fig. S2B**); likely due to this, they exhibited directional mispredictions by the distilled AlphaGenome model less frequently than other variant classes (**Fig. 2B**). Even among these already prominent drivers, we again observed that directional mispredictions coincided with weaker predicted effects (**Fig. 2A**) that often flipped back in other model folds (**Fig. 2B**). Similarly, genes with negatively correlated personal genome predictions had less prominent drivers (**Fig. 2C**) and usually reverted back to positive when predictions were drawn from a different model replicate (**Fig. 2D**).

Together, our results suggest that most causal variants have not only underestimated effect sizes that fail to prominently stand out against background SNVs, but also noisy (unstable over model replicates) underlying attributions. These weak and noisy attributions lead to wrong directional variant predictions by chance, suggesting confident directional mispredictions are a rare failure mode, and the more concerning phenomenon is causal variant underestimation. This is consistent with past reports that wrong directional variant effect predictions do not fall inside coherent motif attributions – further suggesting they reflect noisy, low-confidence signal [11] – and reports that past-generation S2F models also exhibit unstable eQTL SNV predictions [18]. Similarly, without confident directional mispredictions to hold the ranking of individuals in the wrong direction, we observe that negative cross-individual correlations are also likely to be chance occurrences. More broadly, cross-individual correlations routinely flip over zero between model replicates, reflecting the pervasiveness of causal variant underestimation via the absence of highly prominent causal variants with stable attributions to anchor the personal genome predictions in the correct direction against background SNVs with unstable attributions (see **Fig. S5** for examples of anchoring). These results further suggest causal variant underestimation is an important issue affecting most loci, and the common cause underlying important pre-existing modeling challenges.

### 2.3 Exploring the magnitude of causal variant underestimation, its causes, and when predictions can be trusted

To better understand precisely how much more attention state-of-the-art S2F models must pay to causal SNVs for adequate fine-mapping, we reconstructed linearized personal genome predictions with artificially increased or decreased fm-eQTL SNV effect magnitudes (**Methods**). We also flipped the variant effect prediction sign of these SNVs if they were not supported by the data, because our results suggest that stronger, more prominent attributions will coincide with less incorrect variant directional effects (**Fig. 2A**). Holding attributions for background SNVs fixed, we report that an average increase of at least 50-100x in causal variant prediction magnitudes would mostly rescue personal genome prediction accuracy (**Fig. 3A**). Still, there is wide variation due to differences in pre-existing fm-eQTL SNV prominence values, with the weakest group of fm-eQTL SNVs (15% of total) requiring a 1000x increase instead.

**Figure 3:**
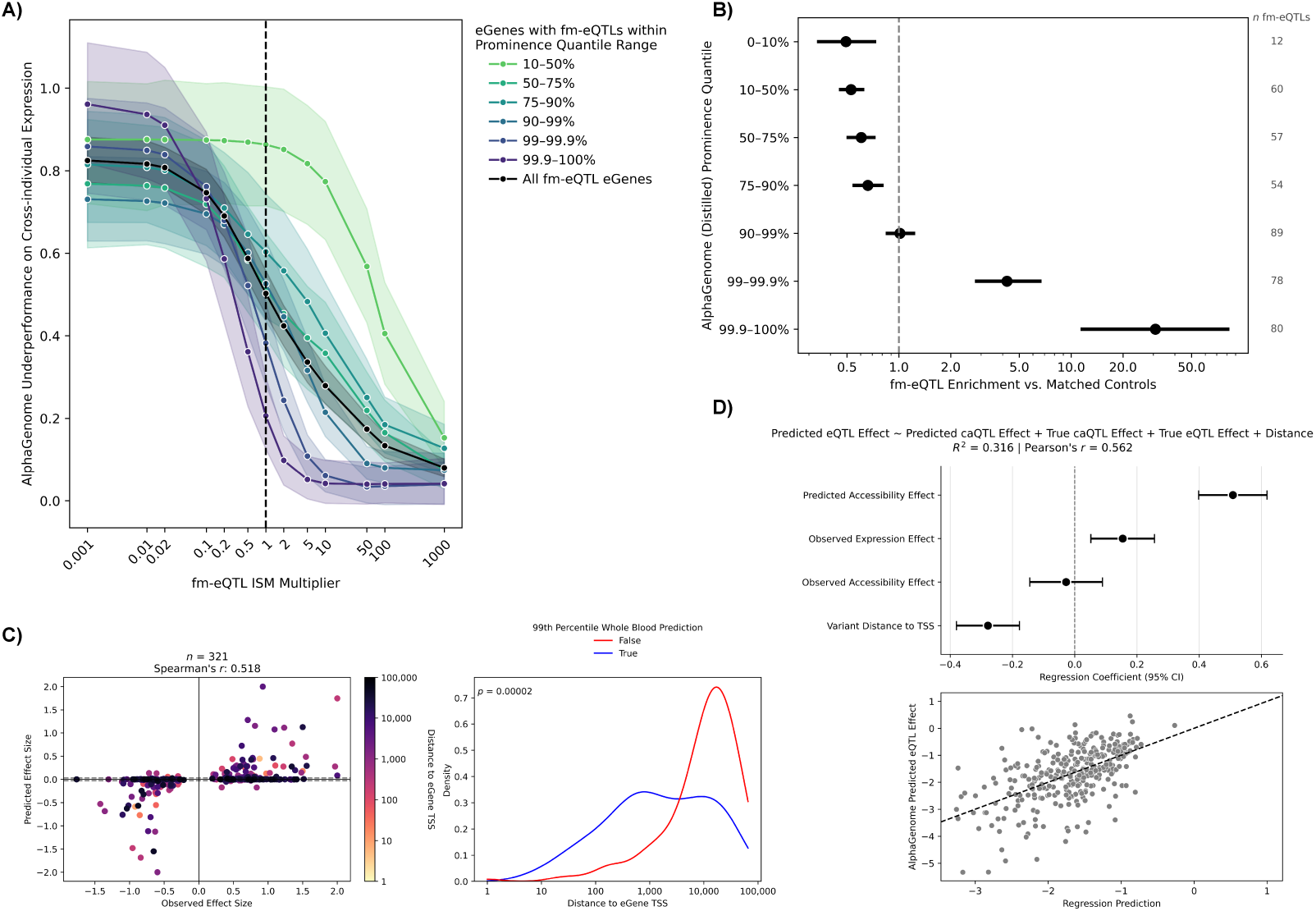
The magnitude of causal variant underestimation, its contributing factors, and when variant effect predictions can be trusted. **A)** fm-eQTLs stratified by prominence range (color), and AlphaGenome’s underperformance on the personal genome task for eGenes linked to fm-eQTLs in each prominence bin (Y-axis) is plotted versus the scaling factor applied to each fm-eQTL before reconstructing personal genome predictions (**Methods**). All groups combined are shown in black, revealing an average 50-100x increase in effect sizes is needed to mostly rescue personal genome predictions, but this increases to 1000x for the least prominent fm-eQTLs. When multiple fm-eQTLs are linked to the same eGene, their prominence is averaged for binning. **B)** fm-eQTLs stratified by prominence range (Y-axis) and their enrichment in these prominence ranges relative to controls matched by TSS distance and MAF. Controls cannot be in LD with fm-eQTLs. The number of fm-eQTLs sampled in each bin divided by the number of controls is the enrichment statistic, and this was repeated after resampling controls 10,000x. Error bars are 95% confidence intervals over resamples. **C)** Predicted and observed effect sizes of MPM-eSNVs colored by TSS distance. SNVs with weaker predicted effects tend to also be further, with exceptions. **D)** Multiple linear regression of AlphaGenome’s predicted eQTL effect magnitude on its predicted caQTL effect magnitude, the measured caQTL and eQTL effects, and the variant’s distance to TSS, using 304 LCL fm-eQTLs that fall within an ATAC-seq peak and are also called as caQTLs. Regression coefficients (top) show predicted expression effects are associated with predicted accessibility effects, independent of distance and true expression effects. AlphaGenome predicted expression effects versus regression model predicted expression effects using these covariates (bottom).

Although we report that AlphaGenome pervasively underestimates causal SNVs, we also observed that the SNVs with the most prominent attributions were the most likely to be causal. Indeed, SNVs with the most prominent, 99.9th percentile prominence predictions were highly enriched for fm-eQTLs (**Fig. 3B**) and wrong directional effect predictions were rare, even at lower MAFs where effect estimates are less reliable (**Fig. 2A**). Further, this threshold is also approximately where predicted variant effects stabilize across model replicates (**Fig. 2A**) and personal genome predictions become highly accurate (**Fig. 1F**) – likely because causal SNVs are upweighted prominently above background SNVs (**Fig. 1G**), indicating effective fine-mapping. We therefore conclude that AlphaGenome has a low overestimation rate – akin to a low false positive rate in classification settings – and predictions can be considered more trustworthy and actionable when predicted effect sizes are simply very large (within the top *∼*0.1% prominence genome-wide). Due to the substantial underestimation rate, however, most causal SNVs fall beneath this range, where their predicted effects are weak, appear indistinguishable from those of gene-matched background SNVs, and flip direction by chance.

Having established that causal variant underestimation is an overlooked, yet pervasive issue responsible for personal genome, mispredicted directional effect, and fine-mapping issues, we next investigated the causes. Underestimated MPM-eSNVs were significantly further from a target gene’s transcription start site (TSS) on average (**Fig. 3C**), consistent with the well-documented neglect of distal regulatory elements by S2F models [19]. Yet we also observed cases where fm-eQTLs are underestimated despite sitting in close proximity to the TSS (**Fig. 3C**).

To better understand the cause, we turned to a set of 304 GEUVADIS lymphoblastoid cell line (LCL) fm-eQTLs [20] that were also inside of a GEUVADIS LCL ATAC-seq peak and were also called as chromatin accessibility QTLs (caQTLs) [21, 22]. As we expect these fm-eQTLs-caQTLs cause expression changes by way of altering chromatin accessibility, we used this dataset to assess the extent to which upstream issues with model detection of variant-induced chromatin accessibility changes lead to weaker downstream predicted expression effects. The magnitude of AlphaGenome’s predicted accessibility effects had a significant positive effect (*p* = 1.3 *×* 10*^−^*^17^ from linear regression) on the magnitude of its predicted expression effect for the same SNVs, independent of distance and the true expression and accessibility effects of these QTL SNVs (**Fig. 3D**). Thus, weaker model detection of variant effects onto local enhancer activity may independently act as a bottleneck for S2F expression predictions.

## 3 Discussion

We showed that causal variant underestimation is a major overlooked issue in S2F expression prediction modeling, directly hurting fine-mapping applications in most loci. Our results indicate the most trustworthy, actionable predictions are also the strongest (top *∼*0.1% prominence quantile), while remaining predictions are generally unstable and miscalibrated. Other reported S2F modeling issues, including wrong directional variant effect predictions and negative cross-individual correlations [11, 12], are chance observations that coincide with high model uncertainty when variant effect attributions are weak. We speculate that attempts to complement statistical fine-mapping with S2F model predictions [8, 9] will improve with stronger predictions for underestimated causal variants.

Identifying underestimation as the underlying issue, we then identified two contributing failure modes: a failure to detect variants’ local effects on enhancer activity, and — even where those effects are detected — inadequate linking of distal elements to their target genes. Our findings unify and extend existing challenges facing S2F gene expression predictions, while clarifying the path forward: rather than being symptoms of different underlying issues, the neglect of distal regulatory elements by these models helps explain their poor personal genome predictions, unstable directional variant effects, and fine-mapping errors, by causing variant underestimation. Thus, to improve S2F models the field must focus on their linking of distal elements to their target genes, thereby yielding stronger attributions for distal variants that would help resolve each of these issues at their source. Independent of distance, underestimated variant effect predictions of local regulatory element activity may act as an upstream bottleneck, also causing expression underestimation; identifying the cause may enable further improvements of expression predictions.

As many groups have attempted, but ultimately failed, to meaningfully improve S2F gene expression predictions by fine-tuning S2F models on paired whole genome sequences and gene expression values [14, 15, 23], we speculate that observational human genetic variation data is insufficient to meaningfully rescue these failure modes. Instead, perturbational datasets (e.g., massively parallel reporter assays [24], CRISPRi-FlowFISH [25]) that sample a larger DNA sequence space may be necessary.

Our results also explain how S2F models can paradoxically show strong aggregate performance on fm-eQTL benchmarks while dramatically underperforming simple linear baselines on personal genome prediction tasks: these tasks are not equally sensitive to underestimation and fine-mapping capabilities. Strong performance on standard variant effect benchmarks only require relatively few well-predicted causal variants with high leverage and is agnostic to attributions for non-causal background SNVs in the same loci, whereas personal genome tasks are highly sensitive to low prominence and compare explicitly against these background SNVs. This discrepancy motivates the use of more challenging and translatable evaluations that directly compare variant effect predictions to others from the same locus, like personal genome ones, that are sensitive to the underestimation failure mode.

Our study has several important limitations. First, we only analyzed fm-eQTLs and not other types of causal variants, such as causal rare variants, which are more challenging to identify and weakly contribute to cross-individual expression differences. Issues concerning the reproducibility of fm-eQTLs across cohorts may also suggest overconfidence in their causal status in some cases [26, 9], raising the possibility that AlphaGenome disagrees with statistical fine-mapping rather than underestimating causal variants. We therefore minimized our reliance on fm-eQTLs and strongly deferred to AlphaGenome’s own attributions, using MPM-eSNVs and personal genome predictions to assess whether it upweights any plausible source of the eQTL association above putatively non-causal background SNVs, and found it does not in most cases. Second, our analysis focused exclusively on AlphaGenome, although we expect our results extend to the entire class of S2F models, which reportedly underperform AlphaGenome on local and distal variant effect tasks that underlie the underestimation issue. Indeed, we repeated a subset of key experiments with Enformer [2] and found that our results are not specific to the choice of S2F model, which is supported by observations of underestimation from various S2F models in the literature [3, 9]. Third, we depend on AlphaGenome’s four cross-validation folds to measure uncertainty in the distilled model. Using the coarse uncertainty range from these four folds, we found evidence to support the claim that weaker variant effect predictions exhibit higher uncertainty and are generally untrustworthy. There may be instances of stable but weak variant effect predictions we would need more model replicates to identify.

## Acknowledgments and Disclosure of Funding

We thank Nilah Ioannidis, Tony Capra, Sandy Floren, Ruchir Rastogi, Anshul Kundaje, and Sara Mostafavi for discussion and feedback.

## Data availability

All data used in this study comes from publicly available sources, listed in **Methods**. Processed data files to reproduce figures will be made available upon publication.

## Large language model disclosure

We used Claude Opus 5 via a chat interface for writing, code development, and visualization support; all output was reviewed by the authors.

## Ethics approval and consent to participate

No human data was collected in this study. The human genome sequences from GTEx that were analyzed in this study were shared with us through dbGaP data use agreements. The original study obtained ethics approval and participant consent.

## A Methods

### AlphaGenome predictions

We used the AlphaGenome Pytorch implementation [27]. Unless otherwise stated, AlphaGenome predictions come from the distilled (“all_folds”) model, which we expected to have the best performance. For experiments involving cross-validation folds, we used model folds 0-3. Unless otherwise stated, we used 131,072bp DNA sequence inputs, centering them on the gene’s transcription start site, as annotated by GENCODE v49 [28].

To form variant expression effect predictions, we used the “GeneMaskLFCScorer”, with the RNA_SEQ output type, using the gene body mask, 128bp bins, and the “Whole_Blood” gtex_tissue track. We refer to the resulting log fold change scores, which reflect the effect of substituting the alternate allele for the reference allele, as variant effect scores, or in silico mutagenesis (ISM) scores. Positive values correspond to cases where the alternate allele is predicted to increase expression relative to the reference, and vice versa.

We applied the same parameters using the “get_gene_mask” function from AlphaGenome Pytorch to process predictions from each individual’s personal genome sequence, which we obtained from a different study [14]. We had 670 individuals with paired whole genome sequences and Whole Blood RNA-seq from GTEx.

### RNA-seq and eQTL data

We downloaded tissue normalized gene expression data from GTEx v8 [16] (“GTEx_Analysis_v8_eQTL_expression_matrices”) and used these as gene expression values for personal genome predictions and fitting elastic net baselines. We also obtained eQTL betas from GTEx v8, and used the supplied CAVIAR [29] fine-mapping results to identify fm-eQTLs with PIP *>* 0.9. Variants used by GTEx for eQTL mapping include only those with MAF *≥* 1%, so we were unable to consider less common variants in our analysis.

### Elastic net baselines

We fit elastic net models separately for each gene, using ElasticNetCV from scikit-learn [30] and a max_iter value of 2000. Using 530 GTEx training individuals, we fit each elastic net using a matrix of genotypes (0, 1, or 2) for each SNV observed in the same TSS-centered window used to form variant effect and personal genome predictions with AlphaGenome for a given gene. Targets were Whole Blood gene expression values from the same gene in the same individual. See the “RNA-seq and eQTL Data” for more information.

After fitting each model, we formed predictions for a held-out set of 71 GTEx individuals, and we compared these predictions to those from the same individuals in the same sequence window using AlphaGenome.

### Calculation of cross-individual correlation

Using the same 71 individuals from the elastic net test set, we generated a vector of 71 observed gene expression values for each gene. We used individuals from the elastic net test set to ensure the correlations obtained from AlphaGenome and elastic net models were comparable. We then generated a vector of 71 predicted gene expression values (using either AlphaGenome or the elastic net), and took the Pearson correlation between observed and predicted gene expression values for each gene. We define the AlphaGenome underperformance on cross-individual expression as:

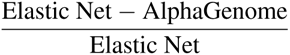

Where Elastic Net and AlphaGenome correspond to their cross-individual correlation value for a given gene. The underperformance value reflects the fraction of elastic net’s cross-individual correlation AlphaGenome does not recover, and we divide by the elastic net value so any underperformance is scaled relative to what the elastic net has shown is achievable for that gene. For example, AlphaGenome achieving a correlation of 60% across individuals while elastic net achieves 80% would suggest AlphaGenome recovers most of the elastic net performance, while an AlphaGenome of 10% and an elastic net value of 30% would suggest the opposite, despite the same absolute difference of 20%.

### Selection of genes for variant effect and personal genome predictions

To focus on genes where genetic variation is expected to contribute to differences in expression between individuals, we first identified genes where an elastic net achieves a cross-individual correlation between predicted and observed Whole Blood gene expression of at least 30%. To conserve computational resources, we first identified these genes using data from a different study [14], where elastic net models were trained on shorter 49,152bp sequences using GTEx data. We then fit elastic nets using 131,072bp sequences on the same genes to verify, and to compare to AlphaGenome’s personal genome predictions. This left us with 713 genes with an elastic net cross-individual correlation of at least 30%, of which 258 have at least one fm-eQTL within the 131,072bp window.

Using these genes, we computed AlphaGenome variant effect predictions for all observed SNVs within the 131kb window. fm-eQTLs, driver SNVs, and MPM-eSNVs, and other SNVs that were around these genes were used for our variant effect prediction analyses. We focused our analyses on fm-eQTLs, MPM-eSNVs, and driver SNVs that could be compared against background SNVs from the same genes, so we did not compute variant effect predictions for SNVs of the same class that come from other genes. Also using these genes, we formed personal genome predictions using AlphaGenome and elastic net models, forming our personal genome dataset.

### Calculation of prominence

Prominence of each SNV is calculated as the log_2_ ratio of its variant effect magnitude divided by the median variant effect magnitude among all SNVs in the window around the same gene.

### Selection of MPM-eSNVs, fm-eQTLs, and driver SNVs

To identify MPM-eSNVs, we used the plink2 [31] r2-unphased calculation, setting a ld-window-kb size of 131, and identifying SNVs with resulting *R*^2^ of at least 90% (LD buddies). We then replaced fm-eQTLs with these SNVs if their variant effect magnitude was greater than that of the fm-eQTL. The resulting set of SNVs (which often include the original fm-eQTLs) were called MPM-eSNVs. We used these to understand if AlphaGenome disagreed with traditional fine-mapping approaches and upweighted different SNVs in LD with fm-eQTLs that could underlie the eQTL association instead. To identify driver SNVs, we implemented a procedure established by previous studies [11, 14]. Briefly, this involves iterating over SNVs around a gene, in descending order of predicted effect magnitude, and identifying the minimum set that can recapitulate AlphaGenome’s personal genome predictions. These SNVs are then said to “drive” the personal genome prediction.

After selecting genes with elastic net cross-individual correlation *>* 0.3 and that had a fm-eQTL within the 131,072bp window, we were left with 258 genes and 366 fm-eQTL/eGene combinations and 321 MPM-eSNV/eGene combinations. There are less MPM-eSNVs because, in cases where there were multiple fm-eQTLs linked to the same eGene and more than one was substituted by the same LD buddy, we dropped the extra instances of this LD buddy so it did not appear more than once. In **Fig. 1E**, we kept only the MPM-eSNV with the strongest ISM magnitude for each gene, leading to 258 points. This was done to avoid situations where there are multiple MPM-eSNVs per gene but not all are underestimated, yet each point for a given gene is associated with the same personal genome metric on the color axis.

In **Fig. 3B**, as we were not comparing variant effect predictions to cross-individual expression predictions from the same genes, we did not require selected genes to have an elastic net correlation *>* 0.3, leaving a larger set of 430 fm-eQTLs (306 unique genes and 419 unique fm-eQTL SNVs).

### Selection of non-eQTLs

To identify non-eQTLs in **Fig. 1G**, we selected SNVs around the same eGene as MPM-eSNVs (and therefore fm-eQTLs) that did not exhibit a significant association with this eGene (determined by looking up “Whole_Blood.v8.signif_variant_gene_pairs.txt.gz” from GTEx [16]). We additionally removed SNVs with MAF *<* 5% to remove less common variants that might not exhibit a significant association with expression due to limited sample size. What remained was a sample of SNVs that we expect are generally sufficiently common to detect an expression association with the same gene, but did not. This served as evidence that these SNVs either do not alter expression of the eGene, or have substantially weaker expression effects than the MPM-eSNV such that these effects are not detectable; comparable predicted effects by AlphaGenome would therefore be considered a fine-mapping error.

### Linear reconstruction of personal genome predictions and artificial modification of fm-eQTL variant effect predictions

To linearly reconstruct personal genome predictions, we took AlphaGenome’s predicted variant effects (ISM scores) for all observed GTEx SNVs within 131kb around a gene across the four cross-validation model replicates. For each individual, we multiplied each of these ISM scores by that individual’s genotype (0, 1 or 2) for that SNV, and summed the products from each SNV together to linearly reconstruct that individual’s personal genome prediction. We repeated this using ISM scores from each of the four model replicates, producing four reconstructed personal genome predictions per individual and gene.

During this procedure, we multiplied fm-eQTLs around each of these genes by a scaling factor. A factor of 1 therefore represents no change, while factors less than one represent artificially weakening the fm-eQTL prediction, and factors greater than one represent artificially strengthening the fm-eQTL prediction. If the fm-eQTL prediction had a predicted directional effect that was inconsistent with its observed effect, we corrected it. We reconstructed personal genome predictions, as specified above, after artificially modifying these fm-eQTLs to assess how modifications to their prominence would alter personal genome prediction accuracy. In **Fig. 3A**, each point represents the median cross-individual correlation over four model replicates, at a given fm-eQTL factor, and error bars represent 95% confidence intervals over genes. We additionally stratified by prominence of the fm-eQTL in the distilled model. If multiple fm-eQTLs were present around one gene, we averaged their prominence before assigning their eGene to one of the prominence bins. We omitted the 0-10% prominence quantile range used in other figures because it included only a single gene after this procedure.

### Comparison of predicted expression effects to predicted accessibility effects using overlapping LCL fm-eQTLs and LCL caQTLs

We obtained GEUVADIS LCL fm-eQTL calls [20] from the eQTL catalog [32] (QTD000110.credible_sets.tsv.gz). We filtered these to keep only SNVs with PIP *>* 0.9 and that sit within AlphaGenome’s 1,048,576bp sequence window. We obtained caQTL calls, also from GEUVADIS LCLs, from [21] (data originally from [22]). We kept caQTL calls that overlapped with fm-eQTL calls and that were inside the same peaks for which they are associated with an accessibility change. Of these, we kept caQTLs with a Benjamini-Hochberg adjusted p *<* 0.05. This left us with 272 unique variants, 299 unique eGenes, and 304 variant-eGene pairs.

To generate AlphaGenome gene expression effect predictions for these variants, we used the GeneMaskLFCScorer, with a gene body mask, 128bp resolution. We used predictions from the “Cells_EBV-transformed_lymphocytes” gtex_tissue track. Sequences were TSS-centered and 1Mb long. To generate accessibility predictions, we used the CenterMaskScorer with a width of 501bp, an aggregation function of diff_log2_sum, 1bp resolution outputs, 1Mb variant-centered sequences, and the GM12878 “atac” track. We verified that various changes to how predictions were formed (e.g., gene body mask vs exon mask, variant-versus TSS-centered sequences, different output tracks) had little impact on final results.

We log_10_ transformed predicted and observed expression and accessibility effects, as well as distance from the eGene TSS, for each of these 304 QTL-eGene pairs. We then fit a linear regression model using statsmodels [33] to predict AlphaGenome’s predicted gene expression value of each QTL SNV, using each SNVs observed expression and accessibility effects, distance to the eGene TSS, and AlphaGenome’s predicted accessibility effect for each SNV as covariates. This enabled us to study the relationship between AlphaGenome’s predicted accessibility and expression predictions, for the same SNV, independent of that SNVs true effects and distance, which would otherwise confound the analysis.

### Enrichment of fm-eQTLs within different prominence quantiles

Starting with 430 Whole Blood fm-eQTLs, larger than other fm-eQTL SNV sets used in this manuscript because we did not further condition on the linked eGene’s gene expression heritability, we first generated a set of control SNVs. For each fm-eQTL, the control SNVs were required to not be in tight LD with the fm-eQTL (do not have *R*^2^ *>* 90%), to have matched MAF that is within 5% as that of the fm-eQTL, and to match their distance to their eGene to be within 1kb of the TSS to that of the matched fm-eQTL relative to its eGene. This led to a median of 1565, and minimum of 501, control SNVs per fm-eQTL. We computed the prominence of each fm-eQTL and control SNV, and separated them into quantile bins. We did not match control SNVs with the fm-eQTL’s eGene because prominence is a gene-controlled metric.

For each fm-eQTL, we sampled one control SNV and counted the number of instances of each group within each prominence quantile bin. We took the fm-eQTL:control ratio of these counts as our enrichment statistic. We repeated this 10,000x and plotted the mean enrichment at each bin, with error bars corresponding to the 2.5 to 97.5 percentile enrichment values among the resamplings.

## B Supplementary figures

**Supplementary Figure 1:**
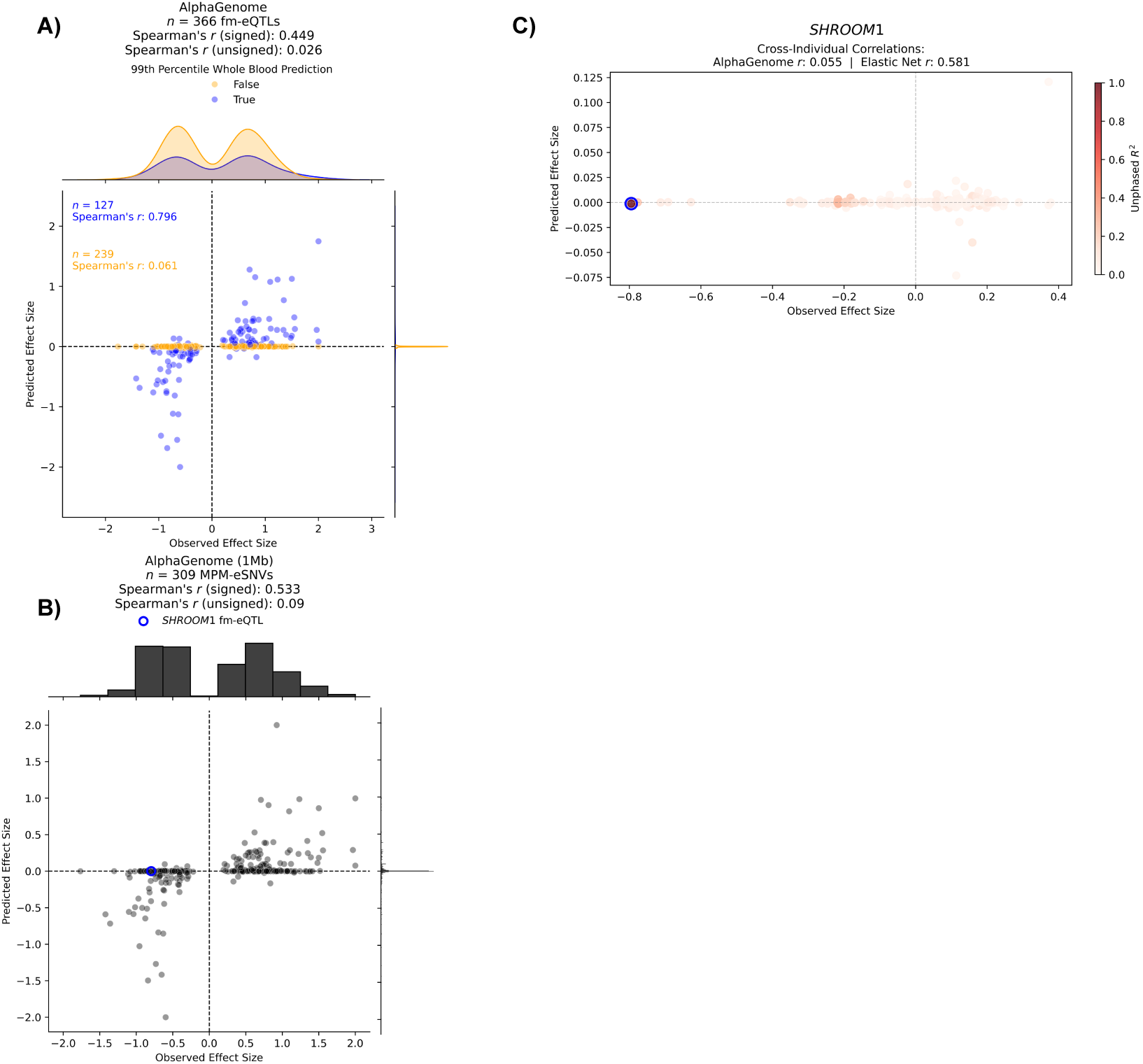
Additional evidence of causal variant underestimation. **A)** Predicted versus observed effect sizes for fm-eQTLs, thresholded (by color) based on whether they are stronger or weaker than the 99th percentile variant effect prediction. Strong aggregate performance on cross-fm-eQTL benchmarks like this one are driven primarily by a subset of well-predicted variants (blue) even while most are underestimated (orange). This is also reflected by comparing the signed versus unsigned correlation coefficient. **B)** Predicted versus observed effect sizes after searching for MPM-eSNVs in 1Mb window. This differs from (Fig. 1A) where MPM-eSNVs were searched for using a 131kb window. The number of variants is lesser here than in (Fig. 1A) because sequence windows were omitted, rather than shifted, if the expansion caused them to extend beyond a chromosomal boundary. **C)** Same as (Fig. 1B) except points are colored by their LD with the fm-eQTL in the region (circled in blue). There are no other upweighted SNVs in tight LD with the fm-eQTL that AlphaGenome is upweighting instead.

**Supplementary Figure 2:**
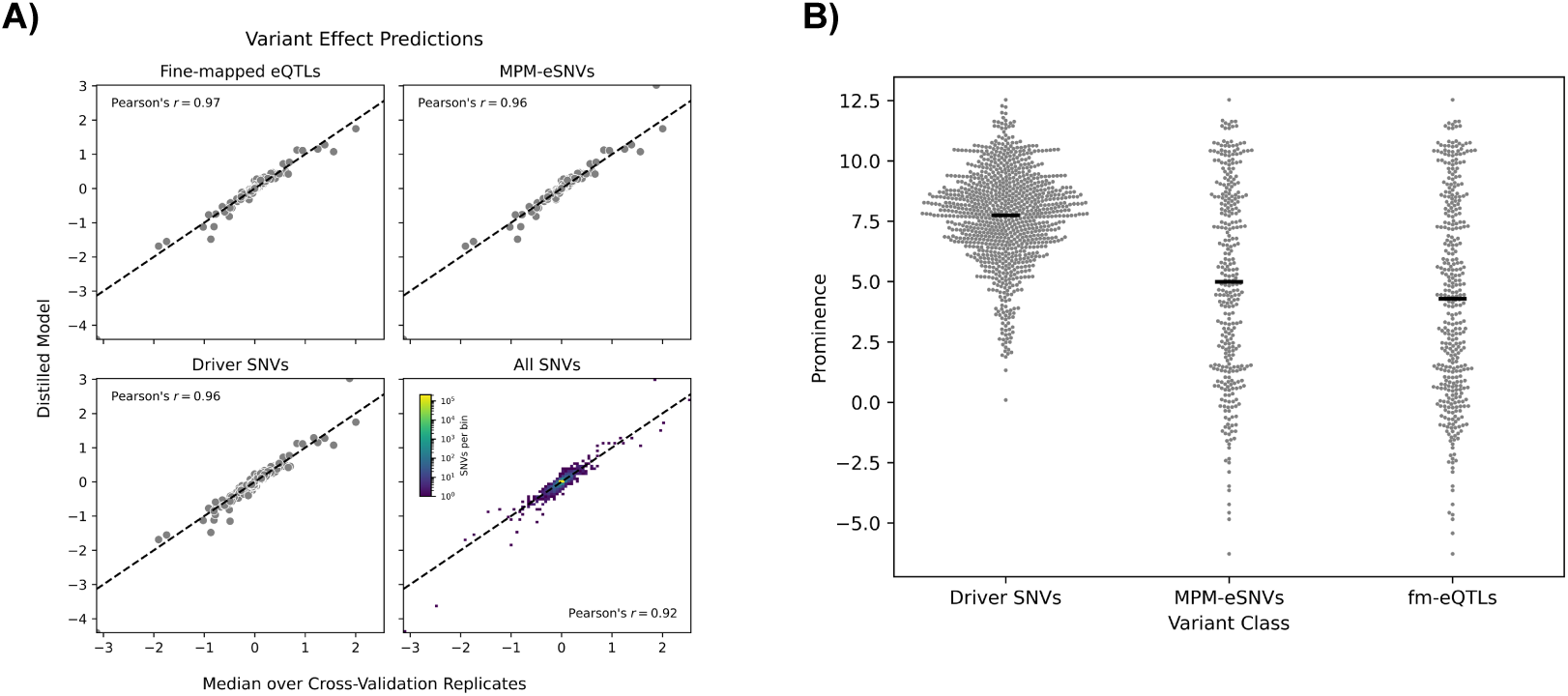
Comparison of predictions over cross-validation folds and variant classes. **A)** Predicted effect sizes for variants from different classes, or all observed SNVs in our dataset (**Methods**, bottom right) from the distilled AlphaGenome model (Y-axis) versus the median over predictions from the four cross-validation folds (X-axis). **B)** Prominence values for variants from different classes.

**Supplementary Figure 3:**
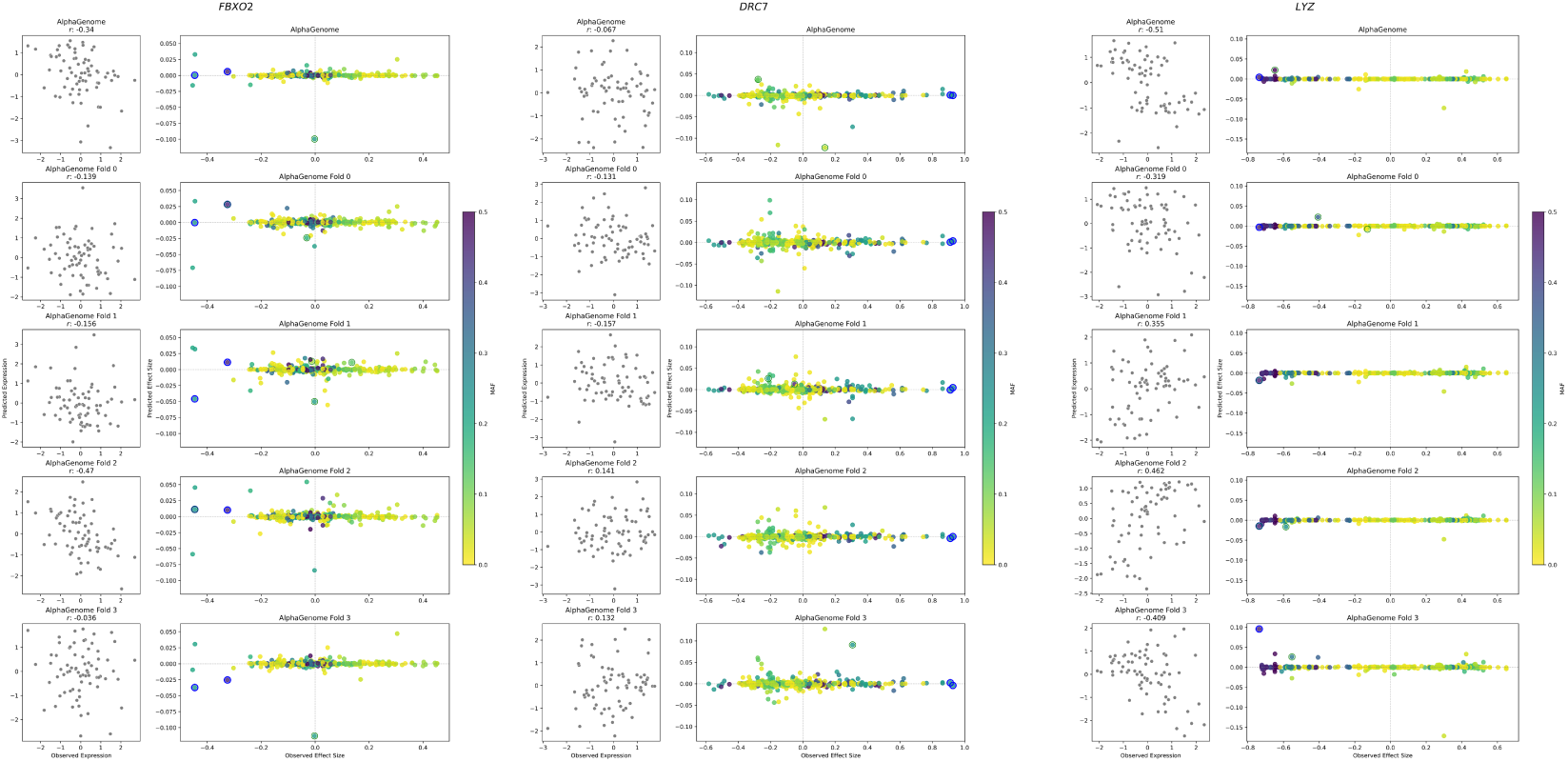
Examples of how insufficiently prominent causal variants lead to inaccurate, noisy personal genome predictions and this reflects poor fine-mapping. Three different genes (columns) for which personal genome predictions are unstable over model replicates (vary substantially from one another and even cross zero) due to lack of sufficiently prominent causal variant attributions. Within each gene’s panel, each row represents predictions coming from either the distilled model (top) or different cross-validation fold replicates. Right column represents predicted versus observed effect sizes for variants around the gene, colored by MAF. fm-eQTLs are annotated in a blue circle, and drivers are annotated in a green circle. Left column represents personal genome predictions for the same gene, reflecting the model’s aggregation of the variant effect predictions on the right, given their genotype. Each point is a different individual and their observed and predicted expression of the focal gene. To accurately fine-map, the model must identify causal variants and prominently upweight them, while downweighting other non-causal SNVs. For each of these genes, causal variants are only weakly upweighted, and have comparable effects to many other SNVs in the window, most or all of which are likely weaker or not functional, indicating poor fine-mapping. Due to the lack of highly prominent SNVs to anchor the predictions, personal genome predictions swing heavily between model replicates due to small changes in variant effect sizes over many variants. This causes cross-individual correlations to cross over zero by chance in different model replicates. For an example of highly prominent attributions that can anchor personal genome predictions and reflect accurate fine-mapping, see **Fig. S5**. Elastic net values for *FBXO2*, *DRC7*, and *LYZ* are 33.6%, 58.2%, and 62.2%, respectively.

**Supplementary Figure 4:**
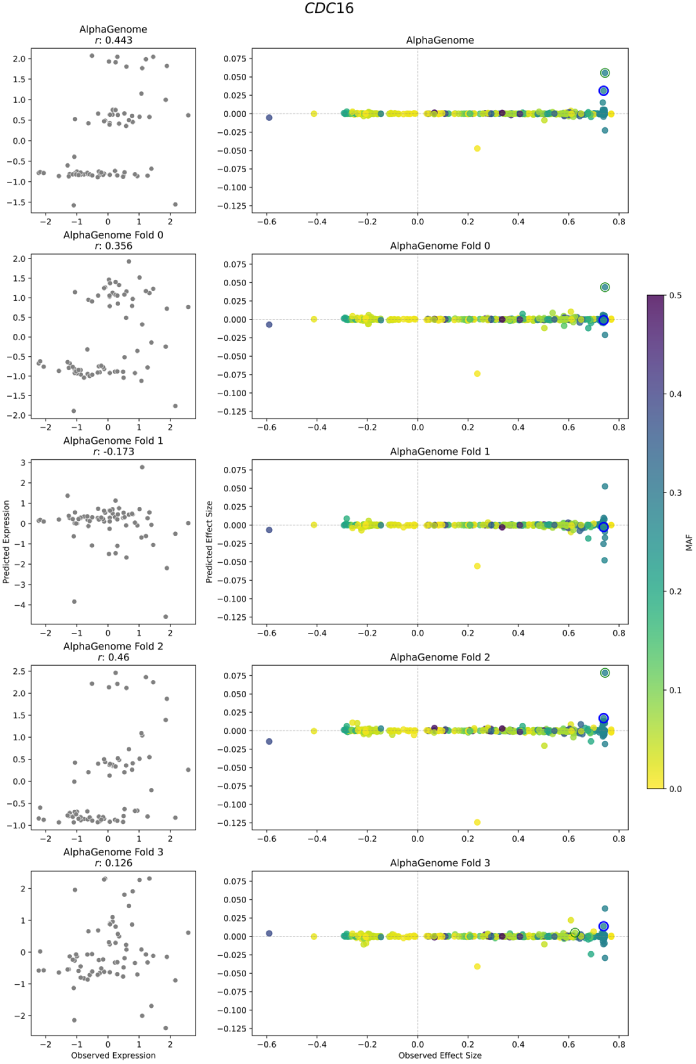
Example of how insufficiently prominent causal variants lead to seemingly accurate personal genome predictions by chance that do not hold in other model replicates. Same setup as **Fig. S3**. Here, fm-eQTL predictions are not highly prominent, so personal genome predictions swing heavily between model replicates, like in **Fig. S3**. Due to random, fortunate combinations of predicted variant effect sizes in some model folds, personal genome predictions appear highly accurate, but this does not hold in other model folds. Thus, causal variant underestimation can lead to seemingly accurate personal genome predictions by chance. Elastic net value for *CDC16* is 50.1%.

**Supplementary Figure 5:**
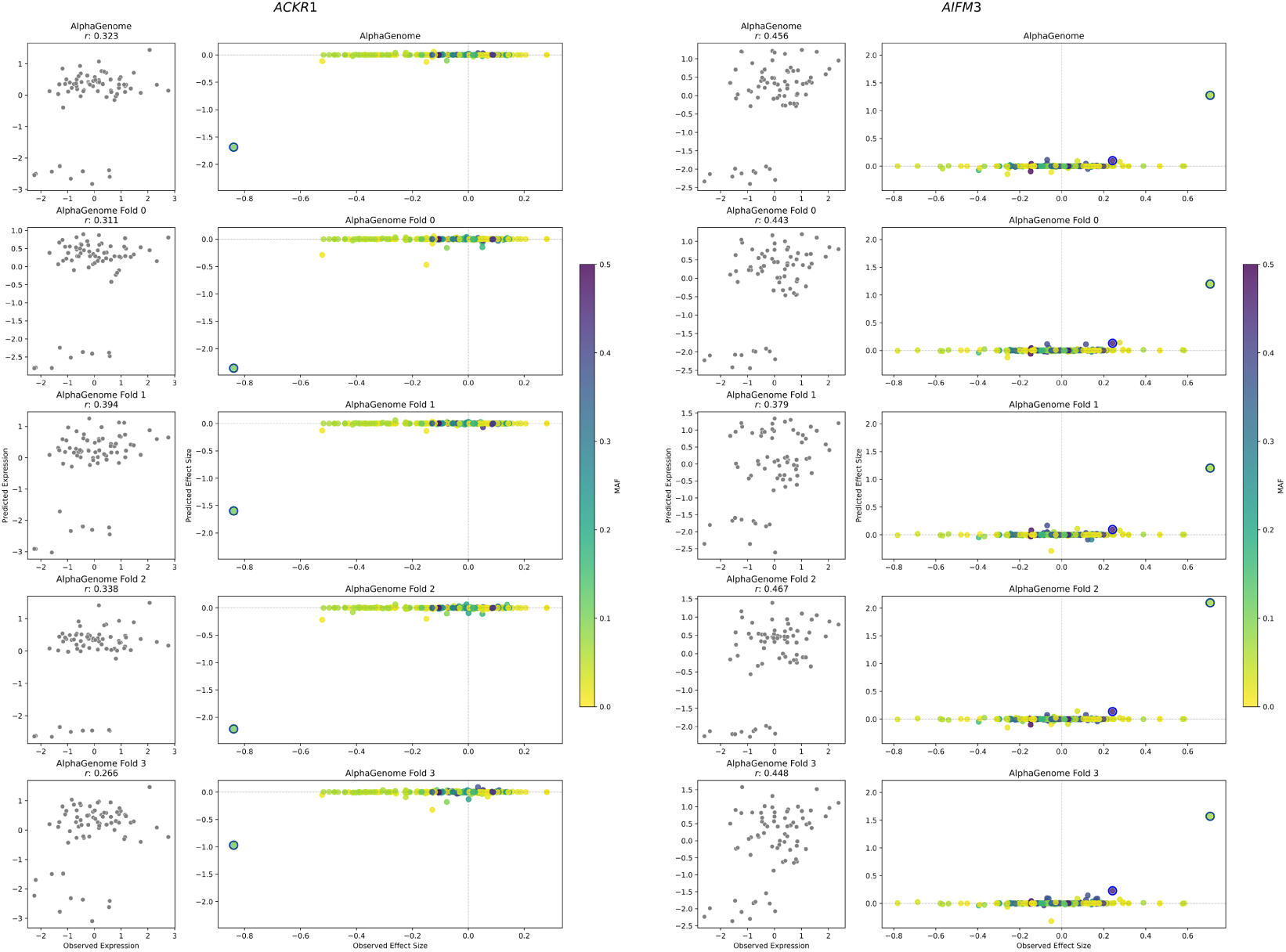
Examples of how highly prominent causal variants lead to accurate, stable personal genome predictions, reflecting successful fine-mapping. Same setup as **Figs. S3-S4**. Here the two genes are selected based on having highly prominent fm-eQTLs, reflecting successful fine-mapping. These prominent fm-eQTLs anchor the personal genome predictions, leading to stable, highly accurate predictions even as variability in variant effect sizes for background SNVs varies between model replicates to the same extent as genes in **Figs. S3-S4**. For *ACKR1*, the elastic net cross-individual correlation is 37.3% and for *PPP1R17* it is 49.5%. In both cases, AlphaGenome achieves scores near these baselines.

**Supplementary Figure 6:**
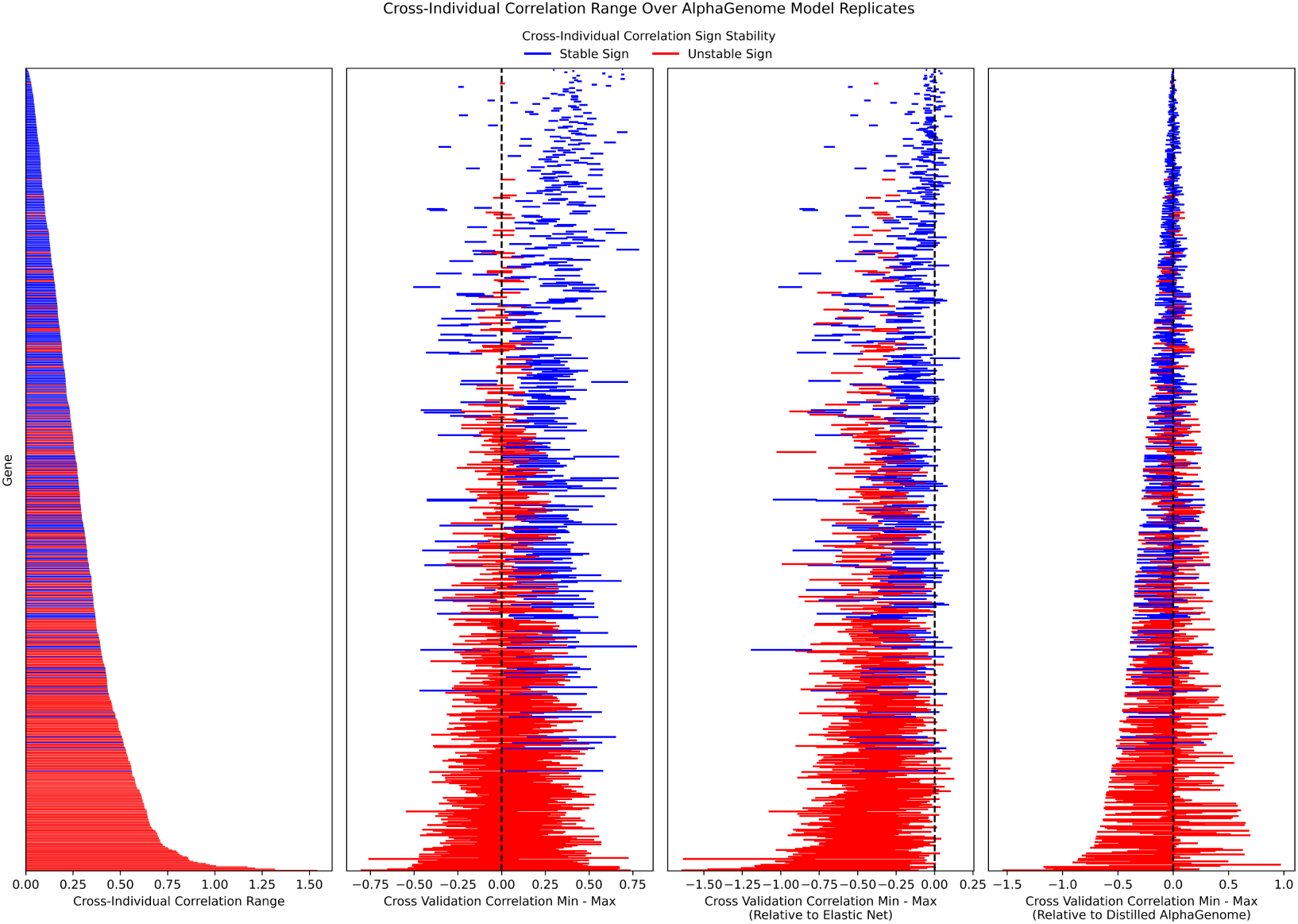
Wide variability in cross-individual prediction accuracy between AlphaGenome model folds. Each row in each panel represents a different gene, ordered the same between all panels. Each bar is colored by whether the cross-individual correlation sign remains the same over AlphaGenome model folds 0-3 (blue), or whether it crosses zero in at least one (red). In the left-most panel, the height of each bar represents the range of cross-individual correlation values found among the cross-validation folds. Most differ substantially, even if the range never crosses zero, likely due to variant underestimation. In the next panel, the actual range between the minimum and maximum value among the four model folds is reported. Many genes that do not cross zero come close and display wide variability nonetheless. We expect many more would cross zero if more model replicates were available. In the next panel, this range is plotted relative to the elastic net baseline value. Negative values reflect distance beneath the elastic net value, while positive values reflect outperformance (and to what extent) of the elastic net. Wide variability leads to similar performance to elastic net models by chance that do not hold in other model folds. The next panel shows this range relative to the cross-individual correlation value achieved by the distilled model. The distilled model typically sits firmly within the distribution from the four cross-validation folds, supporting our usage of them to estimate uncertainty in the distilled model.

**Supplementary Figure 7:**
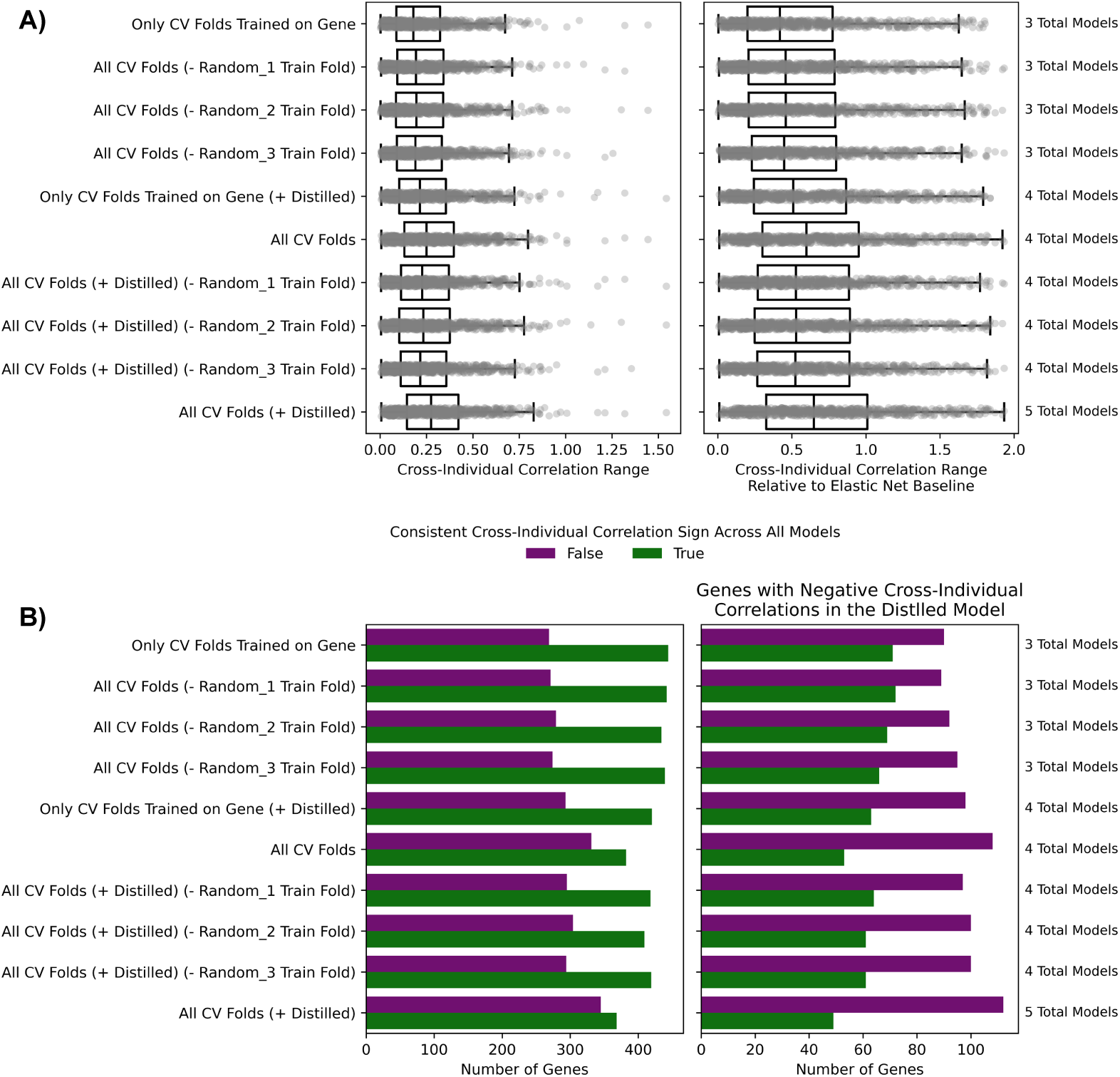
Wide variation in cross-individual correlation range is unrelated to choice of model folds. **A)** Each point is a gene. Range between the minimum and maximum cross-individual correlation value from included models (Y-axis) is displayed on the X-axis (left). The same range, divided by the elastic net value for each gene, is displayed on the X-axis in the panel on the right to show extent of variation relative to a correlation score that has been shown to be achievable for each gene. We observed extensive variation between cross-validation folds (All CV Folds). To ensure this was not caused by highly different, unreliable predictions from the model replicate that did not include a given gene in its train set, we repeated the analysis without it (Only CV Folds Trained on Gene). This reduced variation somewhat, but not by more than if we had kept the omitted fold and removed one of the other three instead (All CV Folds (-Random_Train Fold)), suggesting the reduced, but still substantial, variation comes from decreasing the number of model replicates. Including the distilled model (+ Distilled) led to similar ranges as the cross-validation folds. **B)** Same as top, except counting the number of genes whose cross-individual correlation sign flips at least once (purple) between model folds (among those included on the Y axis). On the right is a subset of these genes that had a negative cross-individual correlation in the distilled model. Similar to in (A), removing the fold that was not trained on each gene led to similar results as removing a random model fold.

